# Neuronal activity in the posterior cingulate cortex signals environmental information and predicts behavioral variability during routine foraging

**DOI:** 10.1101/499814

**Authors:** David L Barack, Michael L Platt

## Abstract

Animals engage in routine behavior in order to efficiently navigate their environments. This routine behavior may be influenced by the state of the environment, such as the location and size of rewards. The neural circuits tracking environmental information and how that information impacts decisions to diverge from routines remains unexplored. To investigate the representation of environmental information during routine foraging, we recorded the activity of single neurons in posterior cingulate cortex (PCC) in monkeys searching through an array of targets in which the location of rewards was unknown. Outside the laboratory, people and animals solve such traveling salesman problems by following routine traplines that connect nearest-neighbor locations. In our task, monkeys also deployed traplining routines, but as the environment became better known, they diverged from them despite the reduction in foraging efficiency. While foraging, PCC neurons tracked environmental information but not reward and predicted variability in the pattern of choices. Together, these findings suggest that PCC mediates the influence of information on variability in choice behavior.

## Introduction

Imagine you are at a horse race, and there are six horses, with Local Field Potential the underdog, facing 100:1 odds against. When LFP wins, a one dollar bet will pay out $100. But in addition to the reward received from this bet, learning that out of the six horses, LFP is the winner reduces your uncertainty about the outcome. Hence, LFP crossing the finish line first yields both reward and information.

Similar problems are often faced by organisms in their environment. Animals must learn not only the sizes of rewards but also their locations, times, or other properties. For example, hummingbirds will adapt their nectar foraging in response to unexpected changes in reward timing (Garrison and Gass 1999). Similarly, monkeys will adapt their foraging routines upon receiving information that a highly valued resource has become available (Menzel 1991). In general, animals can make better decisions by tracking such reward information. Perhaps once a reward has been received, it no longer pays to wait for more because the resource is exhausted or the inter-reward time is too great (McNamara 1982), such as occurs for some foraging animals. Or, perhaps receiving a reward also resolves any remaining uncertainty about an environment (Stephens and Krebs 1986). Keeping track of reward information independent of reward size thus serves as an important input into animals’ decision processes.

We designed an experiment to probe this oft-neglected informational aspect of reward-based decision making. Our experiment is based on the behavior of animals that exploit renewable resources by following a routine foraging path, a strategy known as traplining(Berger-Tal and Bar-David 2015). Trapline foraging has a number of benefits, including reducing the variance of a harvest and thereby attenuating risk(Possingham 1989), efficiently capitalizing on periodically renewing resources(Possingham 1989; Bell 1990; Ohashi, Leslie, and Thomson 2008), and helping adapt to changes in competition(Ohashi, Leslie, and Thomson 2013). Many animals trapline, including bats(Racey and Swift 1985), bees(Manning 1956; Janzen 1971), butterflies(Boggs, Smiley, and Gilbert 1981), hummingbirds(Gill 1988), and an array of primates including rhesus macaques(Menzel 1973), baboons(Noser and Byrne 2010), vervet monkeys(Cramer and Gallistel 1997), and humans(Hui, Fader, and Bradlow 2009). Wild primates foraging for fruit(Menzel 1973; Noser and Byrne 2010), captive primates searching for hidden foods(Gallistel and Cramer 1996; Desrochers et al. 2010), and humans moving through simulated(MacGregor and Chu 2011) and real(Hui, Fader, and Bradlow 2009) environments all use traplining to minimize total distance traveled and thereby maximize resource intake rates.

Though many primates trapline, information about the state of the environment, such as weather(Janmaat, Byrne, and Zuberbühler 2006), the availability of new foods(Menzel 1991), or possible feeding locations(Hemmi and Menzel 1995; Menzel 1996), can influence choices made while foraging. Such detours result in longer search distances and more variable choices(Noser and Byrne 2010; Hui, Fader, and Bradlow 2009) but allow animals to identify new resources(Menzel 1991) and engage in novel behaviors(Noser and Byrne 2010). These benefits are consistent with computer simulations that show stochastic traplining yields better long-term returns than pure traplining by uncovering new resources or more efficient routes(Ohashi and Thomson 2005). Thus environmental information may improve foraging efficiency during routine foraging.

The neural mechanisms that track, update, and regulate the impact of environmental information on decision making remain unknown. Neuroimaging studies have revealed that the posterior cingulate cortex (PCC) is activated by a wide range of cognitive phenomena that involve rewards, including prospection(Benoit, Gilbert, and Burgess 2011), value representation(Kable and Glimcher 2007; Clithero and Rangel 2014), strategy setting(Wan, Cheng, and Tanaka 2015), and cognitive control(Leech et al. 2011). Intracranial recordings in monkeys have found that PCC neurons signal reinforcement learning strategies(Pearson et al. 2009), respond to novel stimuli during conditional visuomotor learning(Heilbronner and Platt 2013), represent value(McCoy et al. 2003), risk(McCoy and Platt 2005), and task switches(Hayden and Platt 2010), and stimulation can induce shifts between options(Hayden et al. 2008). Together, these observations suggest that the PCC mediates the effect of environmental information on variability in routine behavior. However, no studies to date have attempted to separate out the hedonic value from the informational value of rewards in PCC.

Here we tested the hypothesis that PCC preferentially tracks reward information by recording the activity of PCC neurons in monkeys foraging through an array of targets in which environmental information, operationalized as the pattern of rewards, was partially decorrelated from reward size. Monkeys developed traplines in which they moved directly between nearest neighbor targets. As they acquired more information about the state of the environment, their trapline foraging behavior was faster and less variable. While foraging, PCC neurons tracked environmental information but not reward and forecast response speed and variability in traplines. These findings support our hypothesis that PCC mediates the use of information about the state of the environment to regulate adherence to routines in behavior and cognition.

## Materials and Methods

### Task Analysis

Our task manipulates both reward and information. Reward was manipulated by varying the size of received rewards: on every trial, one of six targets had a large reward, one had a small reward that was half of the size of the large, and the remaining four had zero rewards. Information was manipulated by varying the spatiotemporal pattern of rewarding targets. Given a set of four null rewards, one small, and one large, there are 6! distinct permutations. We made the simplifying assumption that monkeys perceived the pattern of received rewards without distinguishing the different null rewards received. This assumption reduces the number of distinct patterns from 720 to 30.

Different patterns correspond to different series of received reward. The environmental entropy *H_E_* contained in receiving some reward (zero, small, or large) depends on the choice number *i* in the sequence and the total number of possible sequences:

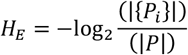

where |·| denotes cardinality, *P* is the set of possible permutations, and {*P_i_*} is the set of remaining permutations after the *i*^th^ choice. The amount of information contained in some reward outcome is computed as the difference in the entropy, what has been learned about the current trial’s pattern of received reward by receiving the most recent outcome:

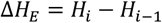

for the amount of environmental entropy *H_E_* on the *i*^th^ outcome. Expected information can then be computed as the mean amount of information to be gained by making the next choice, the weighted average over all possible next information outcomes given the pattern of received rewards thus far:

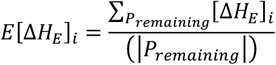

for expected information *E*[Δ*H_E_*] for the *i*^th^ choice, possible outcomes [ΔH_E_]_*i*_ for the remaining permutations *P_remaining_*, and where |·| again denotes cardinality. Thus, as the animal proceeds through the trial, the amount of expected information varies as a function of how many possible patterns of returns have been eliminated thus far. Expected reward *ER* is computed simply as the amount of remaining reward to be harvested on trial *i* divided by the number of remaining targets

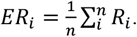

If the animal harvests all of the reward near the beginning of a trial, the expected reward will be zero. However, if the animal does not harvest the rewards until the end of a trial, the expected reward will increase across the duration of the trial.

Given the design of the task, some patterns of returns can have the same amount of expected information for the same choice number but distinct expected rewards (see supplementary information). This partial decorrelation of reward and information occurs as a result of the way expected reward and expected information are computed. Expected information is a function of the number of possible patterns to be excluded by the next outcome, whereas expected reward is a function of the identity of the pattern, and in particular, the amount of reward remaining to be harvested. The same amount of information can be expected from obtaining distinct outcomes because the number of patterns excluded by the different outcomes is identical. But the expected rewards can differ because there are distinct outcomes possible. Hence, our task partially de-confounds information and reward.

### Behavioral Methods

For our behavioral entropy measures, we again used the standard definition of entropy. For behavioral entropy, the probability of a particular step size was computed for each step size by counting the number of trials with that step size and dividing by the total number of trials. Action step sizes (from −2 to 3) and action step size probabilities (probability of taking an action of a given size) were calculated for choices 1 to 2, 2 to 3, 3 to 4, and 4 to 5 (5 to 6 had a constant update of 1). Step sizes were calculated on each choice by determining how many targets around clockwise (positive) or counterclockwise (negative) the next choice was from the previous choice; already selected targets were not included in this calculation. Step size probabilities were calculated by holding fixed all of the covariates for a particular choice (information outcome from previous choice, information expectation for next choice, reward outcome from previous choice, reward expectation for next choice, and choice number) and counting the frequencies for each step size and dividing by the total number of trials with that set of covariates. For each unique combination of covariates (choice number, information outcome, information expectation, reward outcome, and reward expectation), we computed the choicewise behavioral entropy (*H_B_*) for that combination as

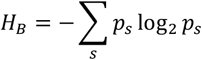

for probability of each step size *p_s_*. Finally, a multilinear regression correlated these behavioral entropy scores with the covariates.

Diverge choices were defined as choices that diverged from the daily dominant pattern (DDP). Determining the DDP relied on assessing the similarity between pairs of trials, for every possible pair on a given day, by computing the pair’s Hamming score(Hamming 1950). To compute the similarity between two trials, each trial’s pattern of choices by target number is first coded as a digit string (e.g., 1, 2, 4, 5, 6, 3). The Hamming distance *D_i,i_*’ between two strings *i, i*’ of equal length is equal to the sum of the number of differences *d* between each entry in the string,

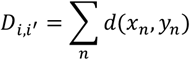

for strings *x, y* of length *n*. We computed *D_i,i_* for every pair of trials, and then, for each unique pattern of choices, computed the average Hamming distance 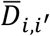. The daily dominant pattern corresponded to the pattern with the minimum 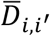.

Response times, defined as the time from end of saccade for the last decision to end of saccade for the current decision, and behavioral entropy were both regressed against a number of variables and their interactions as covariates with multilinear regression. Covariates included choice number in trial, expected information, expected reward, reward outcome from the previous choice, information outcome from the previous choice, and all 2-way interactions. See Table S2 for full results of the response time regression and Table S6 for full results of the behavioral entropy regressions.

### Neural Methods

Two male rhesus macaque monkeys were trained to orient to visual targets for liquid rewards before undergoing surgical procedures to implant a head-restraint post (Crist Instruments) and receive a craniotomy and recording chamber (Crist Instruments) permitting access to PCC. All surgeries were done in accordance with Duke University IACUC approved protocols. The animals were on isoflourane during surgery, received analgesics and prophylactic antibiotics after the surgery, and were permitted a month to heal before any recordings were performed. After recovery, both animals were trained on the traplining task, followed by recordings from BA 23/31 in PCC. MR images were used to locate the relevant anatomical areas and place electrodes. Acute recordings were performed over many months. Approximately one fifth of the recordings were done using FHC (FHC, Inc., Bangor, ME) single contact electrodes and four fifths performed using Plexon (Plexon, Inc., Dallas, TX) 8-contact axial array U-probes in monkey L. No statistically significant differences in the proportion of task relevant cells were detected between the populations recorded with the two types of electrodes (χ^2^, p > 0.5). All recordings in monkey R were done using the U-probes. Recordings were performed using Plexon neural recording systems. All single contact units were sorted online and then re-sorted offline with Plexon offline sorter. All axial units were sorted offline with Plexon offline sorter.

Neural responses often show non-linearities(Dayan and Abbott 2001), which can be captured using a generalized linear model(Aljadeff et al. 2016). We used a generalized linear model (GLM) with a log-linear link function and Poisson distributed noise estimated from the data to analyze our neuronal recordings, effectively modeling neuronal responses as an exponential function of a linear combination of the input variables. We analyzed the neural data in two epochs: a 500 ms anticipation epoch, encompassing a 250 ms pre-saccade period and the 250 ms hold fixation period to register a choice, and more focally the 250 ms pre-saccade epoch itself. Covariates included choice number in the trial, expected information, expected reward, information outcome from the last choice, reward outcome from the last choice, and all 2-way interactions.

In addition to this GLM, we confirmed our model fits in two ways for each neuron: first, we plotted the residuals against the covariates, to check for higher-order structure, and second, we used elastic net regression, to check that our significant covariates were selected by the best-fit elastic net model(Zou and Hastie 2005). Plotting residuals revealed no significant higher-order structure. Furthermore, elastic net regression confirmed our original GLM results (see supplemental methods). None of the significant covariates identified by the original GLM received a coefficient of 0 from the elastic net regression, and the sizes of the significant coefficients identified by the original GLM were very close to the sizes of the coefficients computed by the elastic net regression (see supplemental results).

Perievent time histograms (PETHs) were created by binning spikes in 10 ms bins, time-locked to the event of interest. For the anticipation epoch, PETHs were centered on the end of the choice saccade. For the last informative outcome analysis, PETHs were time-locked to the time of last informative feedback, from two seconds before to two seconds after in 50 ms time bins. PETHs were smoothed with a Gaussian and a 20 ms kernel. To analyze encoding of the last informative feedback, a log-linear GLM regression was then run on vectors of binned spike counts time-locked to the start of the trial, with time in window, time of last informative feedback (a binary covariate encoding whether or not the current time bin was before or after the last informative feedback), and their interaction as covariates.

Step sizes, step size probabilities, and choicewise entropies were linearly regressed against the firing rates during the anticipation epoch, when actions were executed, as reflected in the step size. To assess choicewise entropy encoding before and after receipt of the last bit of information, we calculated the number of neurons that significantly encoded choicewise entropy by choice before the receipt of this information to the number after. For the population response, we first separated trials by mean choicewise entropy across all choices. Next, we compared the normalized average population response for high average choicewise entropy trials to low average entropy during the two seconds before the receipt of the last information. Then we ran the same analysis on the normalized average response during the two seconds following receipt of this information, and reported the results of these two analyses below.

## Results

### Trapline Foraging in a Simulated Environment

To explore the effects of information on divergence from routines, two monkeys (*M. mulatta)* solved a simple traveling salesman problem. In this task, monkeys visually foraged through a set of six targets arranged in a circular array, only moving on to the next trial after sampling every target (Fig. 1B). On each trial, two of the targets were baited, one with a large reward and one with a small reward, with the identity of the baited targets varying from trial to trial. While foraging, monkeys gathered both rewards, herein defined by the amount of juice obtained, and information, herein defined as the reduction in uncertainty about the location of remaining rewards.

**Fig. 1.**
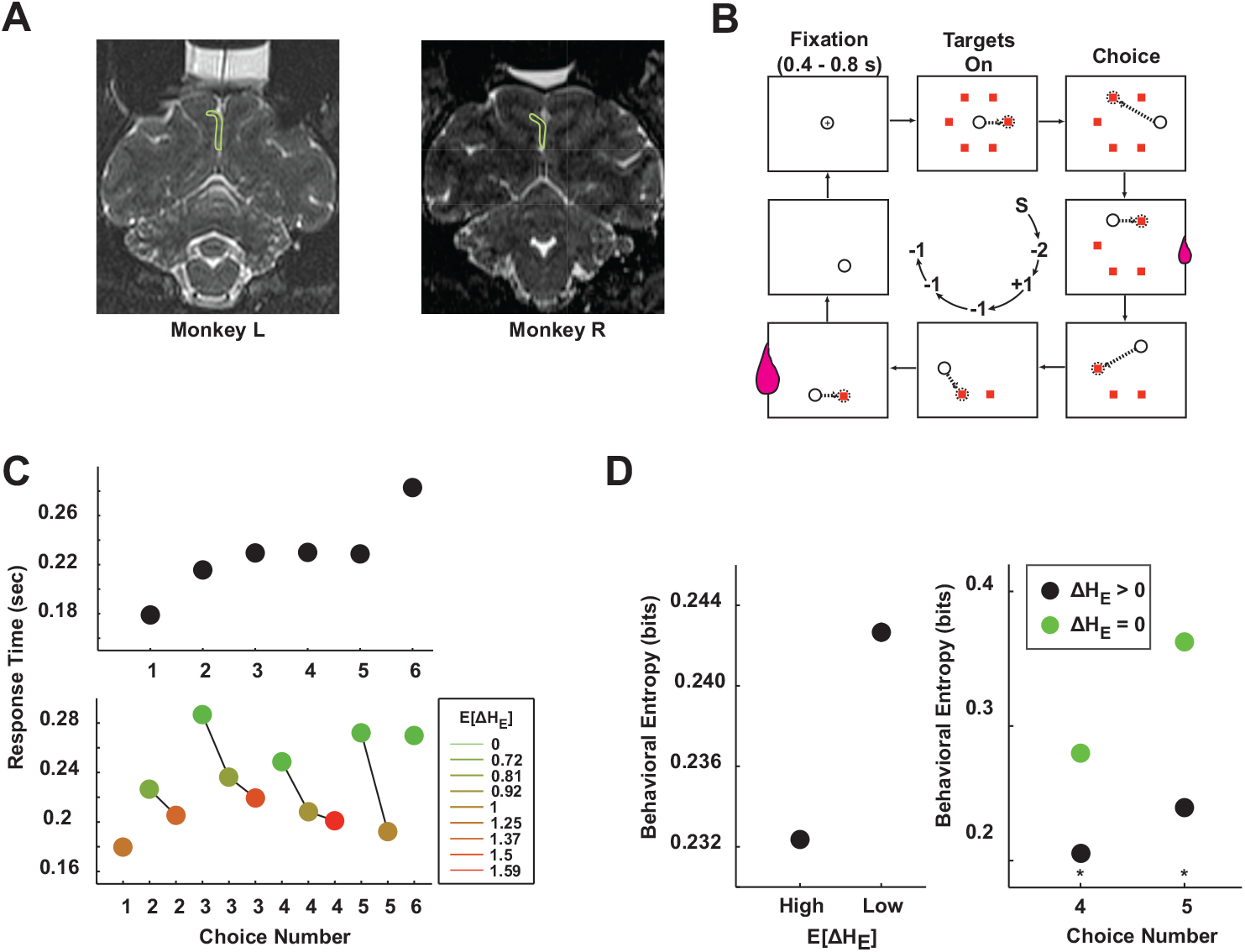
Monkeys spontaneously trapline when foraging in a circular array but diverge from these efficient routines as the environment becomes better known. **A**. Recording location in the posterior cingulate cortex (PCC). Left: Monkey L. Right: Monkey R. **B**. Traplining task, sample trial sequence. Trials began with monkeys fixating a central cross for a variable amount of time. After fixation offset, six targets appeared in the same locations across trials. Monkeys then chose targets in any order. To register a choice, monkeys fixated targets for 250 ms. Only two rewards, one small and one large, were available on every trial, and the identity of the rewarded targets changed in a pseudorandom fashion from trial to trial. In order to advance to the next trial, monkeys had to select every target. Open circle: simulated eye position; dashed arrow: direction of impending saccade; dashed circle: impending saccade endpoint; small juice drop: small reward; large juice drop: large reward. Central circle: step size, the clockwise or counter-clockwise distance between subsequently chosen targets; ‘S’ = start; ‘-2’ = two targets counter-clockwise; ‘-1’ = one target counter-clockwise; ‘+1’ = one target clockwise. C. Top panel: response times increased throughout the trial across all six choices; bottom panel: response times were faster for greater expected information, controlling for choice number in trial. **D**. Left panel: behavioral entropy for high expected information choices (E[ΔH_E_] > mean(E[ΔH_E_])) compared to low expected information choices (E[ΔH_E_] ≤ mean(E[ΔH_E_])); right panel: behavioral entropy for choice numbers four and five, before receipt of last informative outcome (ΔH_E_ > 0; black points) and after (ΔH_E_ = 0; green points). **All** plots: error bars occluded by points; * = p < 0.05, Bonferroni corrected; n = 145,080 choices, 24,180 trials.

By varying the identity of the rewarding targets from trial-to-trial, reward and information were partially decorrelated. Reward was manipulated by varying the size of received rewards, with a small, large, and four zero rewards available on every trial. Information was manipulated by varying the spatiotemporal pattern of rewarding targets. Different patterns correspond to different series of received rewards. Based on the series of rewards received up to a particular choice in the trial, some subset of the set of possible sequences remained, determining the remaining uncertainty for the current trial (see methods). Monkeys gathered rewards and thereby reduced uncertainty about the current trial’s patterns, thereby gathering information about the environment. The amount of information provided by a specific outcome during a trial was calculated as the difference in the entropy before receiving the outcome compared to after. The expected information is a function of the number of possible patterns excluded by the next outcome, whereas expected reward is a function of the remaining reward to be harvested. The expected reward for each target is the total remaining reward to harvest divided by the number of remaining targets. In contrast, the expected information is the mean amount of information to be gained by making the next choice. As the animal proceeds through the trial, the amount of expected information varies as a function of how many possible patterns of returns have been eliminated so far. Distinct possible reward outcomes may offer the same information, and so our task partially de-confounds information and reward. Given the structure of the task, expected reward and expected information are partially decorrelated (linear regression, R^2^ = 0.13).

As monkeys progressed through the array, forecasts of information still available about the foraging environment influenced the speed of responses (all behavioral analyses: *n* = 145,080 choices, 24,180 trials). Though monkeys’ choice response times, the time between target acquisitions, were more sluggish over the course of a trial (Fig. 1C, top panel; linear regression; Both monkeys: *β* = 0.0134 ± 0.0004, p < 1×10^−197^; Monkey L: *β* = 0.0148 ± 0.0005, p < 1×10^−159^; Monkey R: *β* = 0.0100 ± 0.0007, p < 1×10^−39^), the presence of remaining environmental information defined an information boundary generally separating speedy from sluggish choice behavior (Student’s t-test on responses when environmental information remained vs. after; Both monkeys: t(145,078) = −37.88, p < 1×10^−311^; Monkey L: t(103,912) = −38.49, p < 1×10^−321^; Monkey R: t(41,164) = −8.42, p < 1×10^−16^; see supplement and Fig. S1 for full response time regression results). Response times were significantly longer when all information about the array was exhausted than when information remained even after controlling for the number of remaining eye movements (Fig. 1C, bottom panel, response times as function of expected information).

The influence of information on response times suggested a similar influence may hold for the pattern of choices that monkeys made, resulting in trial-to-trial changes in this pattern. Though monkeys usually chose the targets in the same order (the daily dominant pattern, DDP; Monkey R: same DDP across all 14 sessions; Monkey L: same DDP across 24 of 30 sessions; Fig. 1C; see methods), they occasionally diverged from this routine. We found that the informativeness of outcomes influenced this variability in the monkeys’ patterns of choices as measured by behavioral entropy, the expected degree of divergence. First, monkeys’ choices during a trial were egocentrically coded by its step size, the number of targets to the left or right from the current trial’s previously chosen target (Fig. 1B). The probability of a particular step size was computed by counting the number of trials with that step size and dividing by the total number of trials. Anticipation of more informative choice outcomes significantly reduced the entropy of the monkeys’ choices (Student’s t-test; Both monkeys: t(96,718) = −19.25, p < 1×10^−81^; Monkey L: t(69,274) = −3.24, p < 0.005; Monkey R: t(27,442) = −23.99, p < 1×10^−125^; Fig. 1E, left panel; see supplement for full behavioral entropy regression results). While still harvesting information about the current trial, monkeys diverged less from routine traplines, but afterward they diverged more, becoming more variable in their choices (Student’s t-test on choice numbers (CN) 4 or 5; Both monkeys: t(48,358) = −125.98, p ~ 0; Monkey L: t(34,636) = −96.32, p ~ 0; Monkey R: t(13,720) = −71.79, p ~ 0; results also significant for each CN separately; Fig. 1E, right panel). Hence, monkeys diverged less while choices were still informative and more thereafter.

### Environmental Information Signaling by Posterior Cingulate Neurons

We next probed PCC activity during the trapliner task to examine information and reward signaling from 124 cells in two monkeys (Fig. 1A; monkey L = 84 neurons; monkey R = 40 neurons). In order to control for behavioral confounds, all choices where monkeys diverged from traplines were excluded from the analyses in this section (those neural findings are reported in Barack, Chang, and Platt 2017).

Overall, we found that during the anticipation epoch (500 ms encompassing a 250 ms prechoice period and a 250 ms hold fixation period), neurons in PCC preferentially signaled information expectations over reward expectations. An example cell (Fig. 2A) showed a phasic increase in firing rate during the anticipation epoch when expected information was higher for the same choice number in the trial (for example, choice number two (CN2): Student’s t-test, p < 0.0001; firing rate for 0.72 bits = 23.57 ± 1.33 spikes/sec, firing rate for 1.37 bits = 29.79 ± 0.85 spikes/sec). However, after controlling for choice number in the trial and expected information, the same neuron did not differentiate between different amounts of expected reward (Student’s t-test, p > 0.9; firing rate for 0.2 expected reward = 23.73 ± 2.17 spikes/sec, firing rate for 0.4 expected reward = 23.43 ± 1.64 spikes/sec; Fig. 2A, second row from bottom, left panel). The tuning curves for this same cell collapsed across all choice numbers for both expected information and expected reward illustrate the strong sensitivity to large amounts of information (Fig. 2B).

**Fig. 2.**
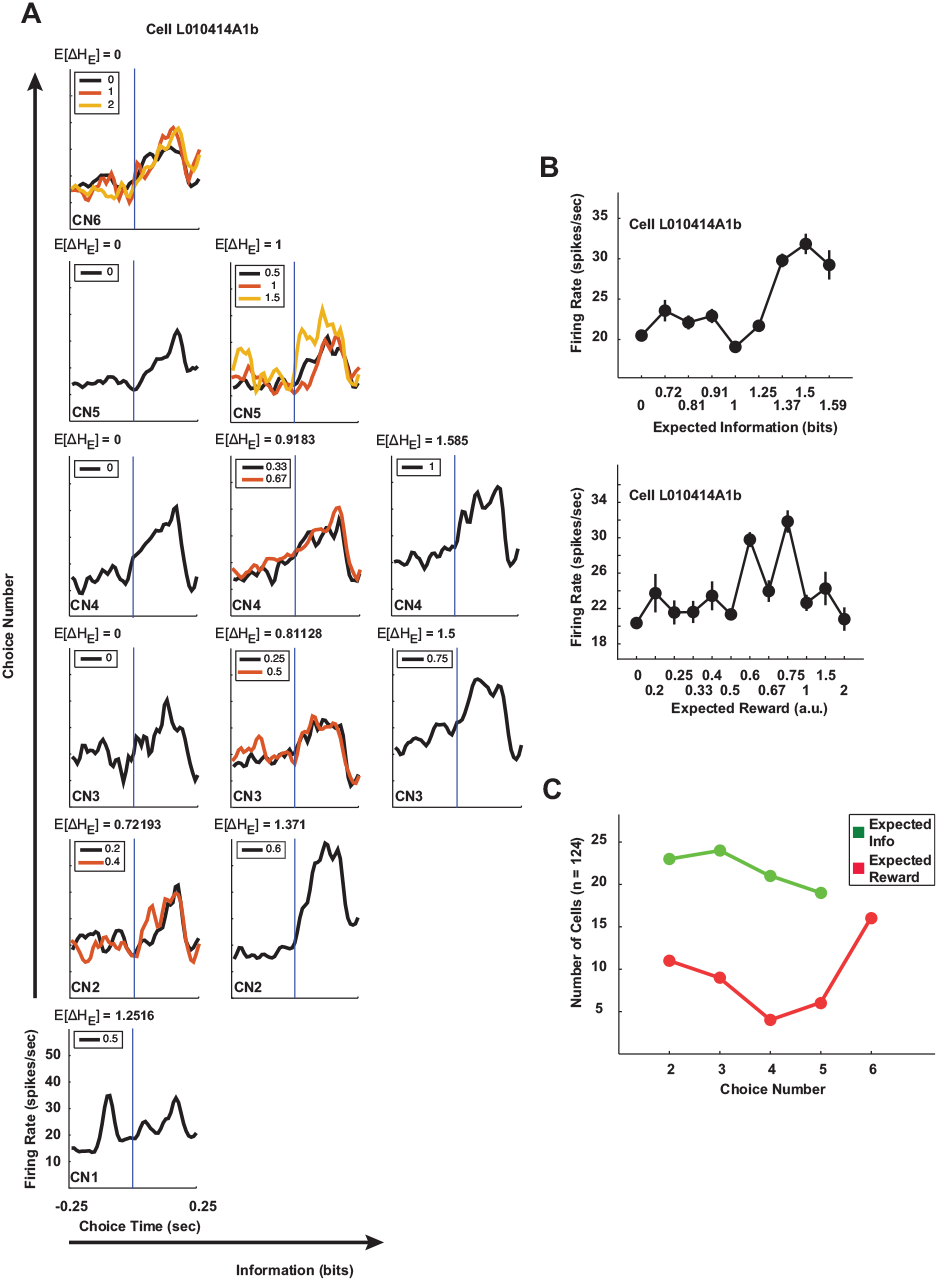
PCC neurons preferentially encode environmental information over reward. **A**. Firing rate of sample neuron encoding expected information but not expected reward across all choice numbers (CN) one through six, plotted separately by expected information (E[ΔH_E_]). Legends indicate expected reward(s) for each plot. Blue line = end of saccade. **B**. Tuning curves for expected information (top panel) and expected reward (bottom panel), collapsed across choice numbers for better visibility, for the cell plotted in **A**. This example cell showed elevated firing rates for higher amounts of information. **C**. Percent of cells encoding expected reward by choice (red) for constant expected information, and percent of cells encoding expected information by choice (green).

In our population of 124 neurons, significantly more cells were tuned to information than reward when controlling for choice number in trial. A generalized linear model (GLM) regression revealed that during the anticipation epoch, 36 (29%) of 124 neurons (Monkey L: 26 (31%) of 84 neurons; Monkey R: 10 (25%) of 40 neurons) signaled the interaction of choice number and expected information, but only 1 (1%) of 124 neurons (Monkey L: 1 (1%) of 84 neurons; Monkey R: 0 (0%) of 40 neurons) signaled the interaction of choice number and expected reward (all results, p < 0.05, Bonferroni corrected; Both monkeys: χ^2^ = 38.9138, p < 1×10^−9^; Monkey L: χ^2^ = 27.5808, p < 5×10^−7^; Monkey R: χ^2^ = 11.4286, p < 0.001; see methods; see supplement for results of full regression and individual monkey results). A direct test for signaling of expected reward, by comparing the average firing rates for different amounts of expected reward for the same choice number and expected information, revealed that only about 10% of neurons signaled expected reward, except on the last choice when all information had been received (Fig. 2C). In contrast, about 20% of neurons signaled expected information (Fig. 2C).

### PCC Neurons Index Response Speed and Variability

We have previously established that PCC neurons signal decisions to diverge from traplines during our task (Barack, Chang, and Platt 2017). However, the extent to which these cells track speed and variability of responses during the task remains to be explored. We next examined the extent to which firing rates in PCC neurons predicted response times across all trials. A high proportion of neurons predicted response times in our task during the pre-saccade epoch, the 250 ms before the end of the choice saccade (both monkeys, 54 (44%) of 124 neurons significantly predicted response times, linear regression, p < 0.05; Monkey L, 46 (55%) of 84 neurons; Monkey R, 14 (35%) of 40 neurons). In order to initially assess the possibility that expected information influenced response times through the PCC, a mediation analysis was run on the group of 9 cells (Monkey L: 9; Monkey R: 2; all variables were z-scored including the firing rates and a variable for cell identity was included in the analysis) that significantly encoded the interaction of expected information and choice number and that significantly predicted response times during that epoch. This analysis compared the direct effect of the interaction of expected information and choice number on saccade response time to the indirect, mediated effect of this variable on response time through PCC neurons (Hayes 2013). Mediation analysis revealed that both paths were small but significant: the direct pathway CN x EI → response times (p < 1 x 10^−6^; estimated effect = −0.0121 ± 0.0024) as well as the indirect pathway CN x EI → firing rates → response times (p < 5 x 10^−10^; estimated effect = −0.0014 ± 0.00022) both significantly sped up responses. Hence, initial indications suggest that PCC partially mediates the influence of expected information on response times.

While PCC neurons predict response speed, we next asked whether these neurons index the degree to which behavior is variable, operationalized as behavioral entropy (see methods). During the anticipation epoch, behavioral entropy varied significantly with firing rate for 70 (56%) neurons (log-linear regression of behavioral entropy against firing rate, p < 0.05; results by choice number: 46 neurons (37%) for CN1, 27 neurons (22%) for CN2, 31 neurons (25%) for CN3, and 41 neurons (33%) for CN4). An example cell was more active for high entropy choices compared to low (Student’s t-test, p < 1 x 10-32; Fig. 3A). Across the population, higher firing rates predicted greater behavioral entropy (Student’s t-test on mean normalized firing rates during anticipation epoch, p < 1×10-5; Fig. 3B). Higher firing rates predicted greater behavioral entropy (BE) in the majority of both the significant subpopulation of 70 cells (GLM, p < 0.05, β_BE_ > 0 in 51 cells, β_BE_ ≤ 0 in 19 cells; mean β_BE_ = 0.0023 ± 0.0011 bitsBE/spike, Student’s t-test against β_BE_ = 0, t(69) = 2.08, p < 0.05) and the whole population (124 neurons; β_BE_ > 0 in 83 cells, β_BE_ ≤ 0 in 41 cells; mean β_BE_ = 0.0058 ± 0.0020 bitsBE/spike, Student’s t-test, t(123) = 2.85, p < 0.01). The percent modulation of behavioral entropy per 1 Hz increase in firing rate can be calculated by dividing the mean regression coefficient for the population by the mean behavioral entropy. In the subpopulation of significant neurons, these mean regression coefficients represent an increase of 2.02% in the average behavioral entropy for every additional spike, and across the whole population, of 1.49% for every additional spike.

**Fig. 3.**
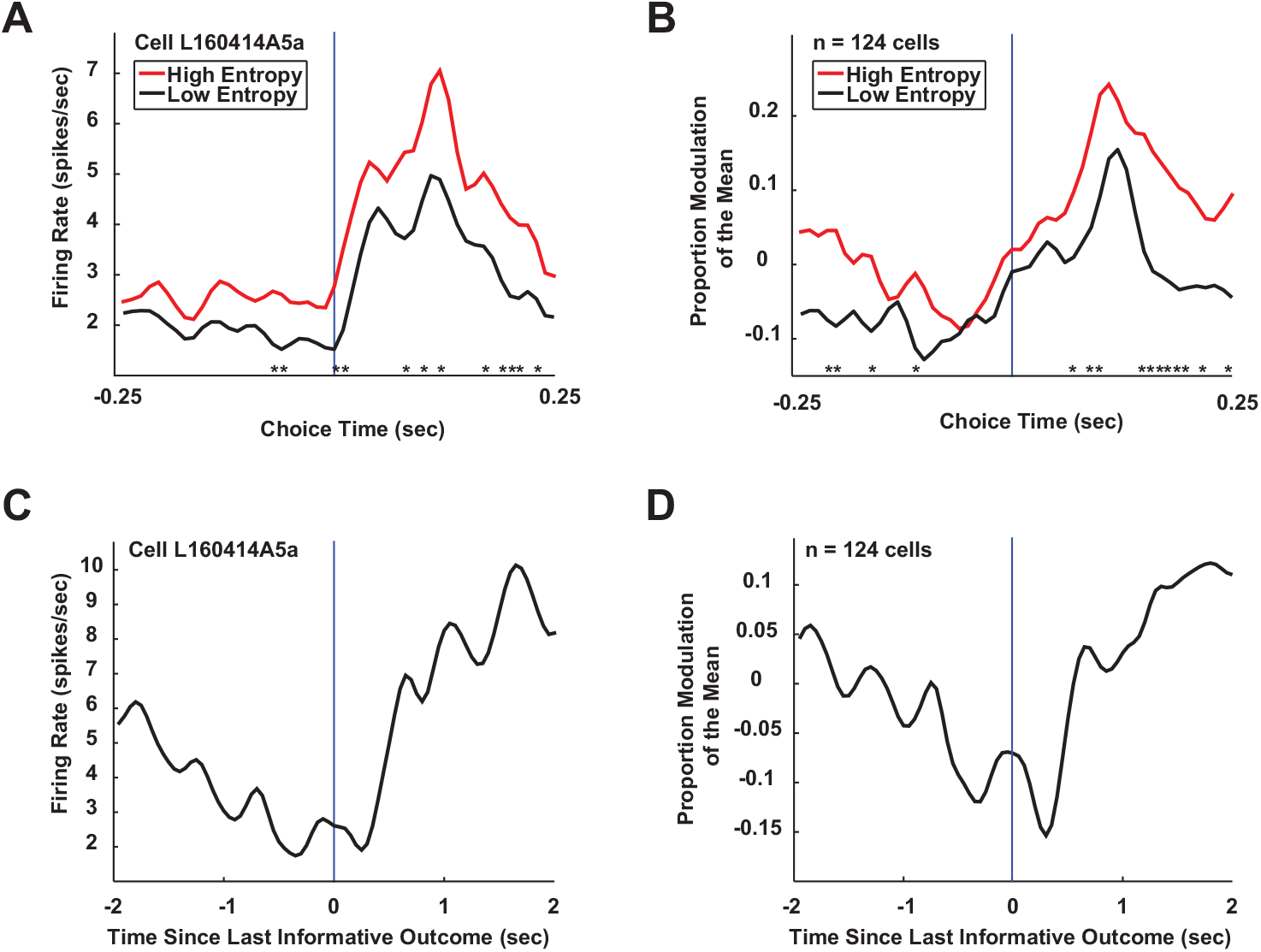
Neurons in PCC forecast divergent behavior. **A**. Same neuron encoding behavioral entropy during the anticipation epoch. This cell was more active for high entropy choices than for low entropy choices. **B**. Population encoding of behavioral entropy. The population was more active for high entropy choices than low. **C**. Sample cell encoding the end of information gathering. This cell had higher firing rates after the last informative outcome compared to before. **D**. The population also encoded this information boundary, with higher firing rates after the last informative outcome compared to before. **B** and **D** plots: n = 125 cells. **All** plots: blue line = end of saccade; * = p < 0.05 for that 10 ms time bin.

Monkeys’ behavior could be characterized as occupying two states along a continuum from routine to divergent. In the more routine state they made faster, less variable choices, while in the more divergent state they made slower, more variable choices. Recall that a boundary defined by the receipt of the last bit of information, when the pattern of rewards on a given trial becomes fully resolved, generally separated periods of routine from divergent behavior. We next investigated whether PCC neurons signaled this information boundary. After a follow-up regression of each trial’s binned spike counts against the time in the trial and the time of last informative outcome, we found that 98 (79%) of 124 neurons differentiated these two states (GLM, effect of interaction, p < 0.05). During a four second epoch centered on the time of the last informative choice outcome, an example cell fired less before that outcome than after (Student’s test, p < 1×10^−56^; Fig. 3C). The population of cells also signaled the transition from routine to divergent behavior, firing more after the last bit of information was received (Student’s t-test, p < 0.005; Fig. 3D).

Finally, behavioral entropy signals and information boundary signals were multiplexed in the PCC population. Significantly fewer cells (χ^2^, p < 1×10^−10^) predicted behavioral entropy after receiving all information (24 (19%) of 124 neurons) than before (74 (60%) neurons). A comparison of PCC population responses on choices with high to low behavioral entropy revealed significantly differences before receipt of the last informative outcome (Student’s t-test, p < 1×10^−4^) but not after (Student’s t-test, p > 0.5; Fig. S2), with greater modulation for high entropy compared to low.

## Discussion

In this study, we show that environmental information influences responses during routine behavior and that the posterior cingulate cortex (PCC) signals this information and predicts response times and behavioral variability. Despite the fact that in our task monkeys could not utilize environmental information to increase their chance of reward, the receipt of environmental information and the exhaustion of uncertainty impacted behavioral routines. Monkeys’ responses were faster and less variable when there was more information to be gathered, but surprisingly slowed and became more variable once the environment became fully known. This pattern of slow responses after resolving all environmental uncertainty departs from the reward rate maximizing strategy of moving in a circle. While monkeys traplined, neurons in PCC robustly signaled information expectations but not reward expectations. These neurons also predicted the speed of responses as monkeys traplined, with neural activity in the subset of the population that signaled information expectations and predicted response times helping to mediate the influence of informativeness on response. PCC also neurons differentiate the degree of behavioral variability before compared to after all information was received about the pattern of rewards, with increasing activity following receipt of the last informative outcome and decreased representations of behavioral variability. In sum, our experimental findings suggest that PCC tracks the state of the environment in order to influence routine behavior.

Monkeys generally chose targets in the same pattern regardless of which targets had been most recently rewarded or informative, consistent with previous findings of repetitive stereotyped foraging in wild primate groups(Noser and Byrne 2007). They also generally moved in a circle, visiting the next nearest neighbor after the current target, likewise consistent with previous findings of in groups of wild foraging primates(Menzel 1973; Garber 1988; Janson 1998). These foraging choices almost always result in straight line routes(Janson 1998; Pochron 2001; Cunningham and Janson 2007; Valero and Byrne 2007) or a series of straight lines(Di Fiore and Suarez 2007; Noser and Byrne 2007). Experiments on captive primates have also observed nearest neighbor or near optimal path finding(Menzel 1973; Gallistel and Cramer 1996; Cramer and Gallistel 1997; MacDonald and Wilkie 1990). Our monkeys’ choices are also consistent with human behavior on traveling salesman problems, wherein next nearest neighbor paths are usually chosen for low numbers of points(Hirtle and Gärling 1992; MacGregor and Ormerod 1996; MacGregor and Chu 2011).

The PCC, a posterior midline cortical region with extensive cortico-cortical connectivity(Heilbronner and Haber 2014) and elevated resting state and off-task metabolic activity(Buckner, Andrews-Hanna, and Schacter 2008), is at the heart of the default mode network (DMN)(Buckner, Andrews-Hanna, and Schacter 2008). The DMN is a cortex-spanning network implicated in divergent cognition including imagination(Schacter et al. 2012), creativity(Kühn et al. 2014), and narration(Wise and Braga 2014). Though implicated in a range of cognitive functions, activity in PCC may be unified by a set of computations related to harvesting information from the environment to regulate routine behavior. Signals in PCC that carry information about environmental decision variables such as value(McCoy et al. 2003), risk(McCoy and Platt 2005), and decision salience(Heilbronner, Hayden, and Platt 2011) may in fact reflect the tracking of information returns from the immediate environment. For example, in a two alternative forced choice task, neurons in PCC preferentially signaled the resolution of a risky choice with a variable reward over the value of choosing a safe choice with a guaranteed reward(McCoy and Platt 2005). Such signals may in fact reflect the information associated with the resolution of uncertainty regarding the risky option. PCC neurons also signal reward-based exploration(Pearson et al. 2009) and microstimulation in PCC can shift monkeys from a preferred option to one they rarely choose(Hayden et al. 2008). Both of these functions may reflect signaling of environmental information as well; for example, the signaling of exploratory choices may reflect the information from an increase in the number of recent sources of reward(Pearson et al. 2009). Evidence from neuroimaging studies in humans similarly reveals PCC activation in a wide range of cognitive processes related to adaptive cognition, including imagination(Benoit, Gilbert, and Burgess 2011), decision making(Kable and Glimcher 2007), and creativity(Beaty et al. 2015). Interestingly, our monkeys adopted a behavioral strategy of slower responses once the state of the environment was fully known. These slower responses may be adaptive insofar as they permit re-evaluation of strategies and potential divergences from routines(Barack, Chang, and Platt 2017).

Uncovering the neural circuits that underlie variability in foraging behavior may provide insight into more complex cognitive functions. A fundamental feature of what we call prospective cognition, thoughts about times, places, and objects beyond the here and now, involves consideration of different ways the world might turn out. Various types of prospective cognition, including imagination, exploration and creativity, impose a tradeoff between engaging well-rehearsed routines and deviating in search of new, potentially better solutions(Gottlieb et al. 2013; Andrews-Hanna, Smallwood, and Spreng 2014; Beaty et al. 2015). For example, creativity involves diverging from typical patterns of thought, such as occurs in generating ideas(Benedek et al. 2014) or crafting novel concepts(Guilford 1959; Barron 1955). During creative episodes the PCC shows increased activity during idea generation(Benedek et al. 2014) and higher connectivity with control networks during idea evaluation(Beaty et al. 2015), perhaps reflecting imagined, anticipated, or predicted variation in the environment. Exploration similarly involves diverging from the familiar, such as in locating novel resources(Ohashi and Thomson 2005) or discovering shorter paths(Sutton and Barto 1998) between known locations. Such prospective cognition requires diverging from routine thought, and the identification of the neural circuits that mediate deviations from motor routines provides initial insight into the computations and mechanisms of prospective cognition. The discovery that the PCC preferentially signals the state of the environment and predicts behavioral variability relative to that state is a first step towards understanding these circuits.

The reinforcement learning literature is replete with models where exploration is driven by the search for information(Schmidhuber 1991; Johnson et al. 2012). These models hypothesize that agents should take actions that maximize the information gleaned from the environment, either by reducing uncertainty about the size of offered rewards(Schmidhuber 1991), the location of rewards in the environment(Johnson et al. 2012), or otherwise maximizing evidence for making subsequent decisions. Furthermore, evidence from initial studies studying information-based exploration shows that humans are avid information-seekers(Miller 1983; Fu and Pirolli 2007) and regulate attentional and valuational computations on the basis of information(Manohar and Husain 2013). In our task, the PCC represented environmental information and tracked when learning about the environment was complete, two variables central to information-based exploration. In particular, the dramatic change in firing rates associated with the end of information gathering suggests that PCC represents the information state of the environment and possibly also the rate of information intake, a central variable in information foraging models(Pirolli and Card 1999; Fu and Pirolli 2007; Pirolli 2007). PCC appears poised to regulate exploration for information.

In sum, harvesting information and behavioral speed and variability were both signaled by PCC neurons, suggesting a central role for PCC in determining how information drives exploration and possibly prospective cognition. Monkeys were sensitive to the amount of uncertainty remaining in the environment, with faster and more reliable patterns of choices while information remained and slower and more variable patterns after environmental uncertainty had been resolved. PCC neurons preferentially tracked this information, and predicted the variability in monkeys’ behavior. Our findings implicate the PCC in the regulation of foraging behavior, and specifically the information-driven divergence from routines.

## Acknowledgments

This work was supported by the National Eye Institute of the National Institutes of Health (R01 EY013496 to M.L.P.) and an Incubator Award from the Duke Institute for Brain Sciences. Correspondence and requests for materials should be addressed to, or to the Center for Science and Society, Fayerweather Hall 511, Columbia University, 1180 Amsterdam Ave., New York, NY 10027.

## Contributions

D.L.B. designed the experiment, D.L.B. collected and analyzed the data, D.L.B. and M.L.P. prepared and revised the manuscript.

## Supplement

To assess the effect of information on behavior independently of reward, we designed a multi-option choice task that partially decorrelated information and reward. Monkeys performed an iterated variant of a Hamiltonian path problem, similar to the uncapacitated traveling purchaser problem, a generalization of the traveling salesman problem(Ramesh 1981; Boctor, Laporte, and Renaud 2003). Hamiltonian path problems are like traveling salesmen problems but the agent is not required to return to the starting location after their trip. In this problem, the agent must efficiently visit a series of markets that may or may not have the desired product, such that in addition to potentially purchasing the product (receiving reward), the agent also learns if the product is available (receiving information). This version of the problem is uncapacitated because the problem assumes that the markets always have sufficient capacity (or supply of a product) to meet the demand. Using this paradigm with varying reward sizes, reward and information can be partially decorrelated.

## Supplemental Methods

### Task Analysis

Our experiment required monkeys to select once each of a set of six targets to harvest the rewards. In every trial in our experiment, two fixed rewards (large and small) were each assigned to one of six locations in the environment in a pseudorandom fashion (Fig. 1B). Trials began with monkeys fixating a central cross for a variable amount of time, ranging from (0.5 – 1 sec). After fixation offset, six targets arranged in a circle appeared. The same locations were used from trial to trial, and monkeys were free to select the targets in any order. To make a choice, monkeys had to fixate their gaze on a target for 250 ms. In order to advance to the next trial, monkeys had to choose each option, even after they’d already harvested the reward available on that trial. Assuming the cost of making a saccade is a monotonic, positive-definite function of distance between targets, the most efficient solution to our task is to minimize saccade times between targets by searching in a circular pattern.

Uncertainty about the current trial’s pattern of received rewards is reduced over the course of the trial as the monkey proceeds through all of the targets. This reduction in uncertainty is quantifiable by examining how many possible patterns are excluded given the rewards revealed by previous choices. For a subset of patterns, the very same information outcome can be delivered by distinct rewards, serving to partially decorrelate and hence de-confound reward and information outcomes. Furthermore, expected reward and expected information, defined as the average amount of information contained in the next outcome given the pattern of rewards received so far, are also partially decorrelated (Table S1).

The linear correlation coefficients between the different task variables (information, expected information, reward, expected reward, etc.) can be computed empirically from the total experienced reward outcomes and information outcomes, and from the total experienced reward expectations and information expectations, derived from the trials the monkeys actually experienced. For the anticipation epoch, this includes expected information and expected reward (R^2^ = 0.1324), expected information and previous choice information outcome (R^2^ = 0.0292), expected information and previous choice reward outcome (R^2^ = 0.1348), expected reward and previous choice information outcome (R^2^ = 2.64 5 8×10^−06^), expected reward and previous choice reward outcomes (R^2^ = 0.1082), previous choice information outcome and previous choice reward outcome (R^2^ = 0.5971). For the outcome epoch, this includes current choice information outcome and current choice reward outcome (R^2^ = 0.3935).

### Neural Regressions

To corroborate our generalized linear model (GLM) regression, we used elastic net regression. Elastic net regression is a model fitting technique that balances two different ways of finding model fits: LASSO (least absolute shrinkage and selection operator) regression and ridge regression. These two model fitting techniques correspond to two ways of computing the distance from the model to the data: using L1 norms (Manhattan distance; LASSO) and L2 norms (Euclidean distance; ridge regression). LASSO regression tends to shift to zero the values of one of two correlated coefficients in order to increase model fit, and ridge regression includes a bias term that helps resolve multi-collinearity in the coefficients. The tradeoff between LASSO and ridge regression is accomplished by means of a parameter α. We used *fmincon,* a constrained search function in MATLAB, to find the maximum likelihood estimate of the best-fit tradeoff parameter on a cell-by-cell, epoch-by-epoch basis (average by epoch: both monkeys: anticipation:, outcome:; monkey L: anticipation: α = 0.9327 ± 0.0132; monkey R: anticipation: α = 0.8963 ± 0.0239). We then compared these fits to the fits from our GLM in three [four?] ways: i. the identity of the covariates that were retained in the best-fit elastic net model; ii. the value of the coefficients computed by the best-fit elastic net model; and iii. the value of just the significant coefficients computed by the best-fit elastic net model. To confirm the identity of the significant covariates, we compared the covariates retained by the elastic net regression to the ones deemed significant by the original GLM. To compare the value of the coefficients computed by the two types of regression, we computed an elastic net score (e_score) for each covariate for each cell for each epoch, equal to the ratio of the elastic net coefficient to the GLM coefficient (i.e., e_score = β_elastic_net_ /β_GLM_). Values around 1 correspond to roughly equivalent computed coefficients; values greater than 1 correspond to a greater weight placed on the covariate by the elastic net compared to the GLM; and values less than 1 correspond to a greater weight placed on the covariate by the GLM compared to the elastic net. Finally, we also computed e_scores but for the significant covariates only.

**Table S1:** Equations for expected reward, entropy, information, and expected information for the reward sequence.

| Patter<br>n<br># | Permutati<br>on<br>$P$ | Expected Reward<br>$ER_i = \frac{1}{n} \sum_i^n R_i$ | Entropy<br>$H_i = -\log_2 \frac{(\vert \{P_i\}\vert )}{(\vert P\vert )}$ | Information<br>$I_i = H_i - H_{i-1}$ | Expected Information<br>$EI_i = \frac{\sum_{P_{remaining}} I_i}{(\vert P_{remaining}\vert )}$ |
| --- | --- | --- | --- | --- | --- |
| 1 | 0 0 0 0 1<br>2 | 0.5 0.6 0.75 1.0 1.5<br>2.0 | 0.5850 1.3219 3.3219 3.9069 4.9069<br>4.9069 | 0.5850 0.7370 1 1.5850<br>1 0 | 1.2516 0.7370 1 1.5850<br>1 0 |
| 2 | 0 0 0 0 2<br>1 | 0.5 0.6 0.75 1.0 1.5<br>1.0 | 0.5850 1.3219 3.3219 3.9069 4.9069<br>4.9069 | 0.5850 0.7370 1 1.5850<br>1 0 | 1.2516 0.7370 1 1.5850<br>1 0 |
| 3 | 0 0 0 1 0<br>2 | 0.5 0.6 0.75 1.0 1.0<br>2.0 | 0.5850 1.3219 3.3219 3.9069 4.9069<br>4.9069 | 0.5850 0.7370 1 1.5850<br>1 0 | 1.2516 0.7370 1 1.5850<br>1 0 |
| ⋮ | ⋮ | ⋮ | ⋮ | ⋮ | ⋮ |
| 30 | 2 1 0 0 0<br>0 | 0.5 0.2 0 0 0<br>0 | 2.5850 4.9069 4.9069 4.9069 4.9069<br>4.9069 | 2.5850 2.3219 0 0 0<br>0 | 1.2516 2.3219 0 0 0<br>0 |

- 1 Each column in the table (except for the leftmost) contains six columns, each corresponding to a choice number during the trial. - 2 Total number of choices on every trial = 6. Key: $|\cdot|$ = cardinality of · ; R = reward; n = choice number.

### Additional Behavioral Results: Response Times

Response times grew longer over the course of the trial (linear regression, p < 1×10^−197^; Fig. 1D, top panel), consistent with past findings that saccade response grow more sluggish over the course of a sequence(Zingale and Kowler 1987). Despite this increasing sluggishness, monkeys were faster to select targets when expected information for making the next choice was higher (multilinear regression, p < 1×10^−9^; Fig. 1D, bottom panel) but surprisingly not when the expected reward for the next choice was greater (multilinear regression, p > 0.93; Fig. S1B). Thus, in addition to increased variability contributing to decreased reward rates, monkeys responses also reflected the state of their knowledge of the environment, growing slower as the trial progressed and further depressing reward rates.

Since different choice numbers in the sequence possess distinct reward and information expectations, response times were regressed against choice number (CN; from 1 to 6), previous information (I-1; from 0 to 2.585 bitsinfo) and reward (R-1; from 0 to 2 unitsjuice) outcomes, expected information (EI; from 0 to 1.585 bitsinfo) and reward (ER; from 0 to 2 unitsjuice), and all 2-way interactions. The regression coefficients, standard errors of the mean, and significance are reported in table S2 and displayed in Figure S1B.

**Table S2:**

| Covariate | Beta | SE | p-value |
| --- | --- | --- | --- |
| Choice number (CN) | 0.0325 | 0.0028 | 0.0000 |
| Expected information (EI) | 0.0236 | 0.0155 | 0.1295 |
| Expected reward (ER) | -0.0833 | 0.0406 | 0.0404 |
| Previous choice information (I-1) | 0.0586 | 0.0109 | 0.0000 |
| Previous choice reward (R-1) | -0.4772 | 0.0496 | 0.0000 |
| CN x EI | -0.0158 | 0.0025 | 0.0000 |
| CN x ER | 0.0005 | 0.0068 | 0.9370 |
| CN x I-1 | -0.0241 | 0.0023 | 0.0000 |
| CN x R-1 | 0.0524 | 0.0062 | 0.0000 |
| EI x ER | 0.0457 | 0.0087 | 0.0000 |
| EI x I-1 | -0.0066 | 0.0108 | 0.5417 |
| EI x R-1 | -0.0092 | 0.0068 | 0.1756 |
| ER x I-1 | 0.0363 | 0.0106 | 0.0006 |
| ER x R-1 | 0.0062 | 0.0046 | 0.1783 |
| I-1 x R-1 | 0.1461 | 0.0136 | 0.0000 |
| Monkey identity | -0.0460 | 0.0017 | 0.0000 |

Like response times after the end of informative outcomes, response times for choices after the animal had harvested the last reward in a trial were significantly longer than for choices when reward remained to be harvested (Student’s t-test, t(145,078) = −46.3694, p < 10^−300^; mean RT for reward remaining = 0.2037 ± 0.00055 sec, mean RT for reward finished = 0.2877 ± 0.0028 sec). Interestingly and surprisingly, we did not see an effect of expected rewards on response times. Naïve regression revealed that response times were faster for higher expected information (linear regression, β = −0.0524 sec/bits, p < 1×10^-317; Fig. S1A, main text) and seemingly for higher expected reward (linear regression, β = −0.0664 sec/units juice, p < 1×10^-282). However, after Bonferroni correction for the number of tests, the only significant covariates that include expected reward also include an information covariate, either expected information (EI) or the previous choice’s information outcome (I-1). In addition, response times were faster following more informative (β_cNxI-i_ = −0.0241 ± 0.0023 sec/bits_info_) but slower following more rewarding (β_cNxR-i_ = 0.0524 ± 0.0062 sec/units_juice_) previous choice outcomes. There was some monkey-to-monkey variability; the influence of the interaction of choice number and expected information was significant in monkey L but, after Bonferroni correction, not in monkey R (tables S4 and S5).

To corroborate the lack of effect of reward size on response times, we directly compared response times for identical trial histories except for the size of past rewards. This test explicitly controls for all other covariates. To directly assess the effect of expected reward on response time, we compared the response times for choices that were matched for the history of choices on that trial but for the size of the rewards, that is, they were matched for the choice number of the choice, information outcomes received thus far, and choice numbers of the rewards received, but not for the size of the rewards received on those choice numbers. Only one such t-test was significant regardless of Bonferroni correction, for response times during the second choice following receipt of a large or small reward on the first choice. Table S3 lists all the p-values from this set of tests. Each row corresponds to the choice number for which response times were compared. Each column corresponds to the choice number in the trial during which either the large or small reward was received, thereby selecting a subset of the response times during the corresponding choice number to compare. For example, the first row (Choice number = 2) picks out all response times during the second choice, and the first column (First choice in trial) picks out the response times for which either a large or small reward was received during the first choice in the trial. The first entry (Choice number = 2, First choice in trial) lists the p-value from the t-test comparing the responses during the second choice where the first choice was the large reward to those response times during the second choice where the first choice was the small reward. As can be observed from Table S3, only the first test is significant.

**Table S3:**
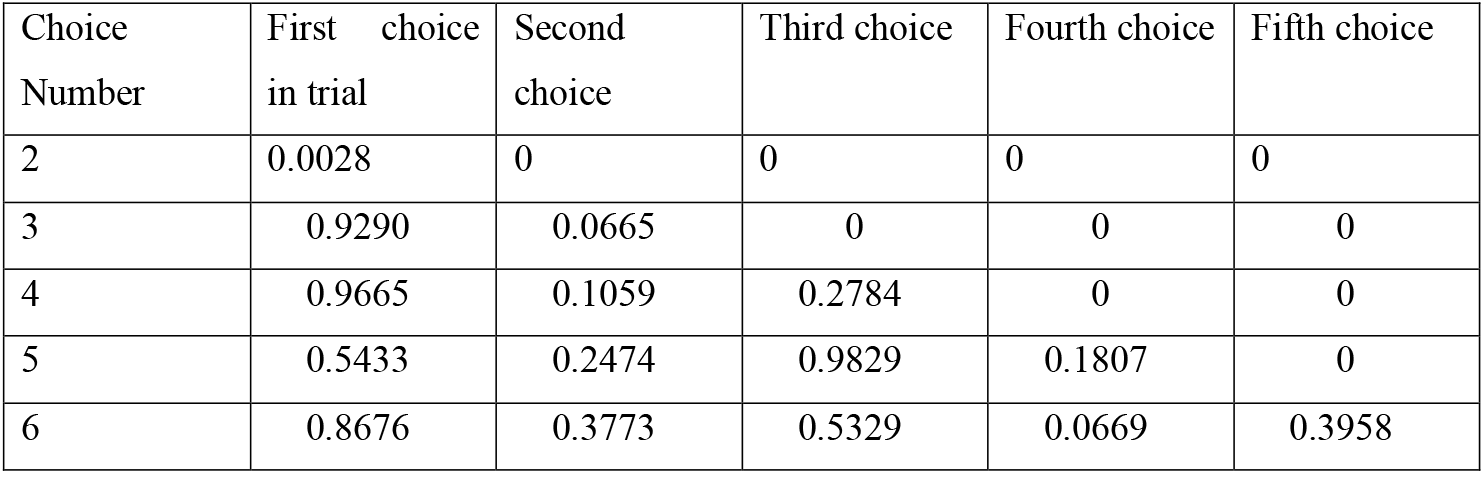

Individual regression results of response times against choice number, expected information, expected reward, previous behavioral information outcomes, previous choice reward outcome, and all 2-way interactions now follow:

**Table S4:**
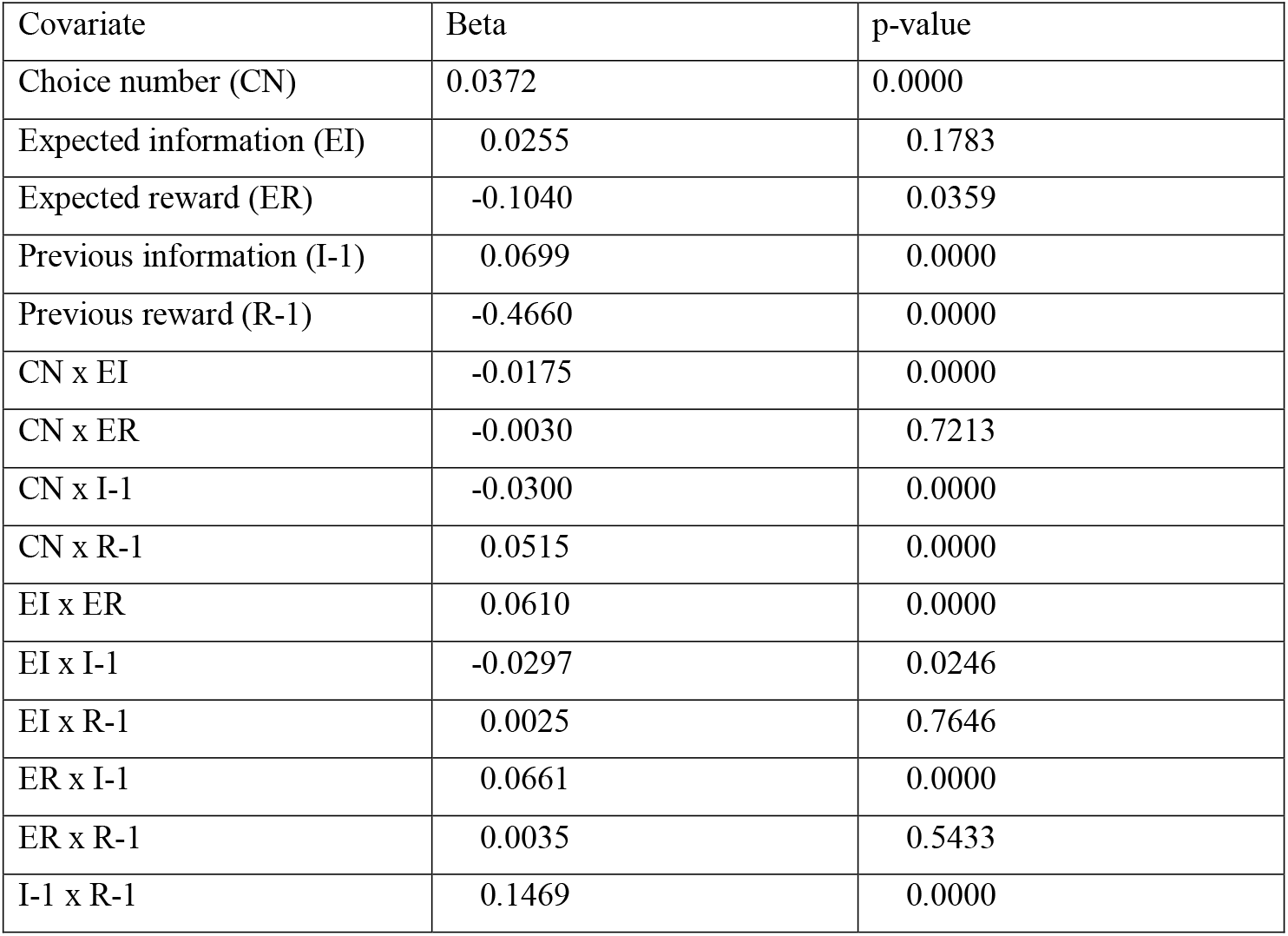
Monkey L (103,914 choices, 17319 trials), RT regression:

| Covariate | Beta | p-value |
| --- | --- | --- |
| Choice number (CN) | 0.0372 | 0.0000 |
| Expected information (EI) | 0.0255 | 0.1783 |
| Expected reward (ER) | -0.1040 | 0.0359 |
| Previous information (I-1) | 0.0699 | 0.0000 |
| Previous reward (R-1) | -0.4660 | 0.0000 |
| CN x EI | -0.0175 | 0.0000 |
| CN x ER | -0.0030 | 0.7213 |
| CN x I-1 | -0.0300 | 0.0000 |
| CN x R-1 | 0.0515 | 0.0000 |
| EI x ER | 0.0610 | 0.0000 |
| EI x I-1 | -0.0297 | 0.0246 |
| EI x R-1 | 0.0025 | 0.7646 |
| ER x I-1 | 0.0661 | 0.0000 |
| ER x R-1 | 0.0035 | 0.5433 |
| I-1 x R-1 | 0.1469 | 0.0000 |

**Table S5:** Monkey R (41,166 choices, 6861 trials), RT regression:

| Covariate | Beta | p-value |
| --- | --- | --- |
| Choice number (CN) | 0.0205 | 0.0000 |
| Expected information (EI) | 0.0170 | 0.5211 |
| Expected reward (ER) | -0.0154 | 0.8228 |
| Previous information (I-1) | 0.0304 | 0.1001 |
| Previous reward (R-1) | -0.4908 | 0.0000 |
| CN x EI | -0.0107 | 0.0108 |
| CN x ER | 0.0070 | 0.5437 |
| CN x I-1 | -0.0088 | 0.0257 |
| CN x R-1 | 0.0532 | 0.0000 |
| EI x ER | 0.0029 | 0.8431 |
| EI x I-1 | 0.0512 | 0.0054 |
| EI x R-1 | -0.0370 | 0.0012 |
| ER x I-1 | -0.0405 | 0.0229 |
| ER x R-1 | 0.0116 | 0.1345 |
| I-1 x R-1 | 0.1392 | 0.0000 |

### Model Fits to the Behavior

Model fits to the behavior revealed that monkeys generally chose targets in the same pattern regardless of which targets had been most recently rewarded or informative.

This finding was corroborated by the daily dominant pattern results reported in the main text. Monkey R chose targets in the same pattern across all 14 recording sessions, whereas monkey L chose targets in the same pattern on 24 sessions, with four patterns accounting for the remaining six days (two days each for two of the patterns and one each for the remaining two patterns).

### Behavioral Entropy Additional Results

From the distribution of step sizes on each choice number, we calculated the behavioral entropy (BE) (0.0294 to 0.5021 bits_BE_; mean_BE_ = 0.1139 ± 0.0170 bits_BE_ across all choices) and correlated these values with choice number (CN), the next choice’s expected information (EI) and expected reward (ER), the previous choice’s information (I-1) and reward (R-1) outcomes, and all 2-way interactions. After combining both monkeys’ data, all covariates significantly influenced behavioral entropy (p < 0.05, Bonferroni corrected) except for the interaction of expected information and reward from the previous choice (EI x R-1, p > 0.6). The effects of previous information outcome (I-1) and expected information (EI) interactions with choice number were both significant across the two monkeys (multilinear regression, all p’s < 0.0000001; both monkeys: β_cNxEI_ = −0.1260 ± 0.0011 bits_BE_/bits_info_; β_cNxI-1_ = −0.0538 ± 0.0008 bits_BE_/bits_info_; Fig. S1C) and significant for both monkeys individually (monkey L: β_cNxI-1_ = −0.0672 ± 0.0009 bits_BE_/bits_info_; β_cNxEI_ = −0.1626 ± 0.0012 bits_BE_/bits_info_; monkey R: β_cNxI-1_ = −0.0181 ± 0.0017 bits_BE_/bits_info_, β_cNxEI_ = −0.0296 ± 0.0024 bits_BE_/bits_info_; Fig. S1c). The effects of the interaction of choice number and reward were significant but inconsistent: later previous reward outcomes resulted in significantly increased variability in the second monkey but not the first, whereas later reward expectations resulted in significantly increased variability in the first monkey but not the second (multilinear regression, all p’s < 0.0000001 except where indicated; both monkeys: β_cNxR-1_ = 0.0235 ± 0.0030 bits_BE_/units_juice_, β_cNxER_ = 0.0775 ± 0.0026 bits_BE_/units_juice_; monkey L: β_cNxR-1_ = 0.0014 ± 0.0033 bits_BE_/units_juice_, p > 0.66, β_cNxER_ = 0.0980 ± 0.0029 bits_BE_/units_juice_; monkey R: β_cNxR-1_ = 0.0734 ± 0.0063 bits_BE_/units_juice_, β_cNxER_ = 0.0064 ± 0.0055 bitsBE/units_juice_, p > 0.24; Fig. S1c). Here we report the full regression results (Table S7).

**Table S6:** Beta coefficient values, standard errors, and p-values from regression of behavioral entropy against the listed covariates and 2-way interactions.

| Covariate | Beta | SE | p-value |
| --- | --- | --- | --- |
| Choice number | 0.1238 | 0.0013 | 0 |
| Previous info<br>outcomes | 0.1882 | 0.0033 | 0 |
| Expected info | 0.4966 | 0.0053 | 0 |
| Previous reward<br>outcomes | -0.1367 | 0.0265 | 0.0000 |
| Expected reward | -0.4285 | 0.0116 | 0.0000 |
| CN x I-1 | -0.0538 | 0.0008 | 0 |
| CN x EI | -0.1260 | 0.0011 | 0 |
| CN x R-1 | 0.0235 | 0.0030 | 0.0000 |
| CN x ER | 0.0775 | 0.0026 | 0.0000 |
| I-1 x EI | -0.0438 | 0.0041 | 0.0000 |
| I-1 x R-1 | 0.0256 | 0.0077 | 0.0009 |
| I-1 x ER | 0.0448 | 0.0039 | 0.0000 |
| EI x R-1 | 0.0013 | 0.0029 | 0.6608 |
| EI x ER | 0.0835 | 0.0025 | 0.0000 |
| R-1 x ER | 0.0907 | 0.0034 | 0.0000 |
| Monkey ID | 0.0110 | 0.0005 | 0.0000 |

We now report the full individual monkey regressions:

**Table S7:**
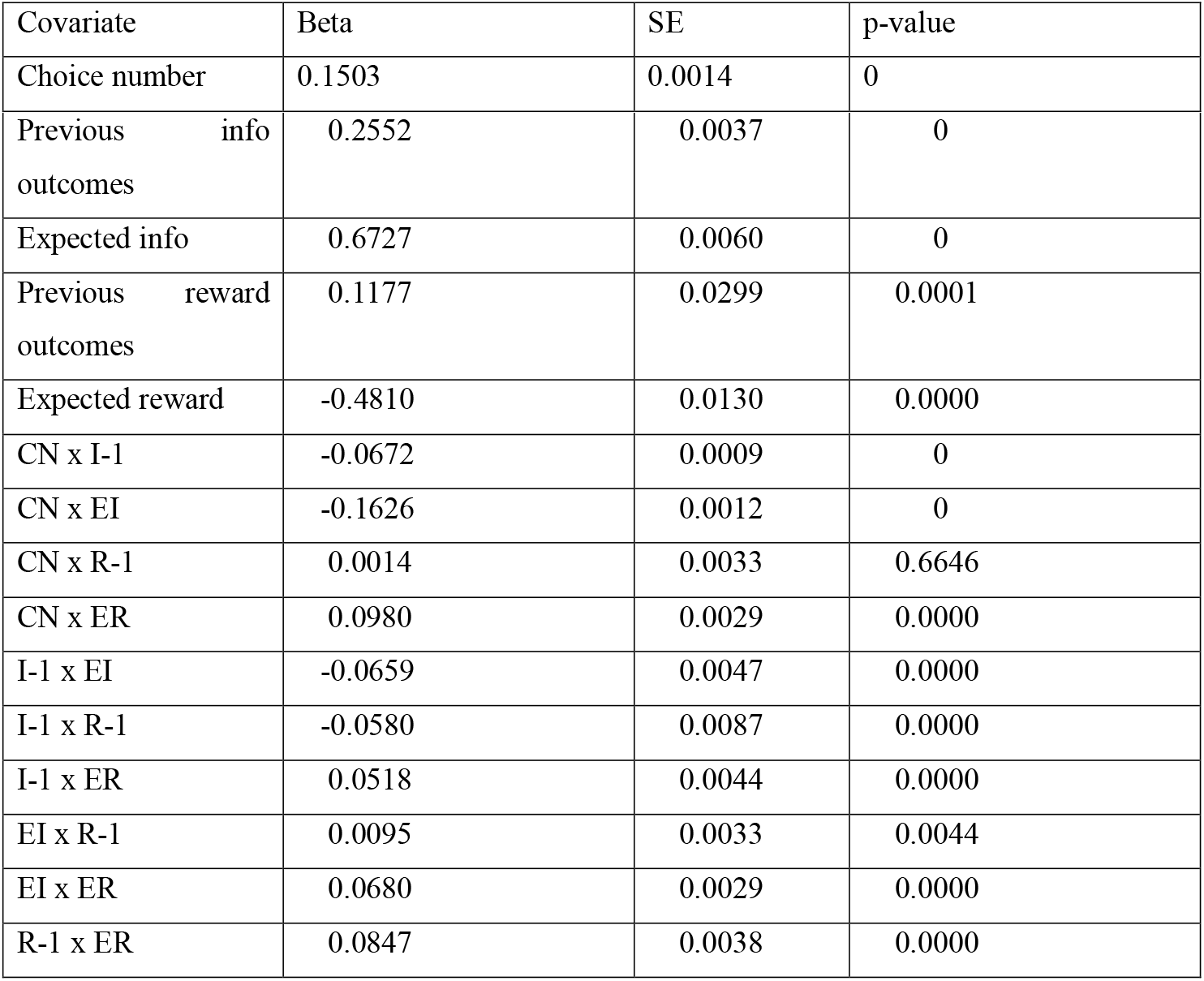
Monkey L (103,914 choices, 17319 trials), behavioral entropy regression:

**Table S8:** Monkey R (41,166 choices, 6861 trials), behavioral entropy regression:

| Covariate | Beta | SE | p-value |
| --- | --- | --- | --- |
| Choice number | 0.0603 | 0.0027 | 0.0000 |
| Previous info<br>outcomes | 0.0196 | 0.0070 | 0.0053 |
| Expected info | 0.0450 | 0.0113 | 0.0001 |
| Previous reward<br>outcomes | -0.7385 | 0.0565 | 0.0000 |
| Expected reward | -0.1875 | 0.0246 | 0.0000 |
| CN x I-1 | -0.0181 | 0.0017 | 0.0000 |
| CN x EI | -0.0296 | 0.0024 | 0.0000 |
| CN x R-1 | 0.0734 | 0.0063 | 0.0000 |
| CN x ER | 0.0064 | 0.0055 | 0.2483 |
| I-1 x EI | 0.0031 | 0.0089 | 0.7224 |
| I-1 x R-1 | 0.2299 | 0.0164 | 0.0000 |
| I-1 x ER | 0.0258 | 0.0083 | 0.0019 |
| EI x R-1 | -0.0159 | 0.0062 | 0.0110 |
| EI x ER | 0.1156 | 0.0054 | 0.0000 |
| R-1 x ER | 0.0993 | 0.0072 | 0.0000 |

The uninformative outcomes could occur for choices four and five, as some choices occurred when the previous choice’s outcome was uninformative, after the current trial’s pattern was already revealed. In the text, we reported that behavioral entropy was greater following uninformative than informative choices, collapsing across choices four and five. Both choices four and five individually reflected this finding (cN_4_: BE_info_ = 0.2050 ± 0.0003678 bits, BE_no_info_ = 0.2801 ± 0.0017 bits, t(24178) = −52.0376, p ~ 0; cN_5_: BE_info_ = 0.2402 ± 0.0005666 bits, BE_no_info_ = 0.3627 ± 0.0012, t(24178) = −96.3793, p ~ 0). The individual monkey results reflected this difference as well (for monkey L: for choice number four: BE_info_ = 0.2907 ± 0.0007 bits, BE_no_info_ = 0.3272 ± 0.0032 bits, t(7857) = −12.4444, p < 1×10^-34; for choice number five: BE_info_ = 0.3481 ± 0.001 bits, BE_no_info_ = 0.4334 ± 0.0017, t(7857) = −37.2945, p < 1×10^-279; for monkey R: for choice number four: BE_info_ = 0.2889 ± 0.001 bits, BE_no_info_ = 0.4264 ± 0.0034 bits, t(6859) = −35.3207, p < 1×10^-250; for choice number five: BE_info_ = 0.3005 ± 0.0012 bits, BE_no_info_ = 0.4455 ± 0.0017, t(6859) = −58.4394, p ~ 0). Thus, the effects were consistent and strong in both monkeys.

### Additional Neural Results: Elastic Net Regression

Our three checks—verifying the presence of the significant covariates, comparing the elastic net weights to the original GLM weights for all covariates, and a similar comparison restricted to only the significant covariate coefficients from the original GLM (see supplemental methods)—confirmed our original GLM analysis. First, none of the significant covariates identified by our original GLM for any cell during any epoch for either monkey received a coefficient of 0 from the elastic net regression. This suggests that our regression results were robust despite the correlations in the covariates.

Second, we compared the e_scores (see supplemental methods) for the elastic net and original GLM regressions. The average e_score = 1.3747 ± 0.5120 (monkey L: e_score = 1.5968 ± 0.7405; monkey R: e_score = 0.9084 ± 0.0823), suggesting that the elastic net regression tended to weight the covariates more than the original GLM, though this was clearly driven by monkey L. (Examination of the individual covariate comparisons further revealed that this effect was driven by the immense weight that the elastic net regression put on expected information for monkey L; see table below.) The following table reports the individual covariate e_scores (mean ± sem). Note that the elastic net regression weighted expected information much more than the original GLM but only for monkey L. All other fits for that epoch were comparatively reasonably close to the original GLM.

**Table S9:** Elastic-net regression results for main GLM.

| Covariate | Monkey L | Monkey R | Both |
| --- | --- | --- | --- |
| Choice number (CN) | $1.1228 \pm 0.2436$ | $1.0377 \pm 0.0728$ | $1.0953 \pm 0.1664$ |
| Expected reward (ER) | $0.8528 \pm 0.0441$ | $0.8867 \pm 0.0337$ | $0.8637 \pm 0.0317$ |
| Expected information (EI) | $11.1105 \pm 10.1277$ | $0.8904 \pm 0.0505$ | $7.8137 \pm 6.8609$ |
| Previous choice info (I-1) | $0.7180 \pm 0.1358$ | $0.6169 \pm 0.1955$ | $0.6854 \pm 0.1112$ |
| Previous choice reward (R-1) | $0.7588 \pm 0.0303$ | $0.7914 \pm 0.0350$ | $0.7693 \pm 0.0234$ |
| CN x ER | $0.8339 \pm 0.0982$ | $0.8451 \pm 0.0446$ | $0.8375 \pm 0.0679$ |
| CN x EI | $0.9729 \pm 0.0595$ | $0.7348 \pm 0.1256$ | $0.8961 \pm 0.0577$ |
| CN x I-1 | $0.9328 \pm 0.0921$ | $0.9516 \pm 0.0424$ | $0.9389 \pm 0.0637$ |
| CN x R-1 | $0.8031 \pm 0.0440$ | $0.8104 \pm 0.0303$ | $0.8054 \pm 0.0313$ |
| ER x EI | $1.0155 \pm 0.0494$ | $0.9859 \pm 0.0530$ | $1.0060 \pm 0.0375$ |
| ER x I-1 | $0.9935 \pm 0.0135$ | $1.0092 \pm 0.0111$ | $0.9986 \pm 0.0098$ |
| ER x R-1 | $1.0699 \pm 0.0803$ | $1.0123 \pm 0.0946$ | $1.0513 \pm 0.0622$ |
| EI x I-1 | $0.9987 \pm 0.0088$ | $1.0003 \pm 0.0300$ | $0.9992 \pm 0.0113$ |
| EI x R-1 | $0.9921 \pm 0.0097$ | $0.9481 \pm 0.0508$ | $0.9779 \pm 0.0176$ |
| I-1 x R-1 | $0.7766 \pm 0.0703$ | $1.1050 \pm 0.3644$ | $0.8825 \pm 0.1266$ |

Third, we ran these same statistics, but restricted only to the significant covariates from our original GLM. The average e_score = 0.9682 ± 0.0035 (monkey L: e_score = 0.9694 ± 0.0039; monkey R: e_score = 0.9615 ± 0.0087), a very good fit between the elastic net results and the original GLM, especially compared to the e_score for all covariates, suggesting that the original GLM was a fair representation of the influence of the different covariates.

### Additional Neural Results: Anticipation Epoch PCC Neuron Responses

In the main text, we reported that PCC neurons preferentially signaled expected information over expected reward. Significantly more neurons encoded expected information than expected reward (χ^2^ = 12.86, p < 0.0005), and significantly more neurons encoded the interaction of choice number and expected information than the interaction of choice number and expected reward (χ^2^ = 23.04, p < 0.000005). This preference was maintained until the last choice number in a trial (CN2: 23 (19%) EI, 10 (8%) ER; CN3: 24 (19%) EI, 9 (7%) ER; CN4: 21 (17%) EI, 4 (3%) ER; CN5: 19 (15%) EI, 6 (5%) ER; CN6: 16 (13%) ER; Fig. 2C). Of the cells significantly encoding the interaction of choice number and expected information, 19 (63%) were negatively encoding and 11 (37%) were positively encoding, with a mean regression weight (β_CNxEI_) = −0.11 ± 0.075 (spikes/sec)/(choice*bitsinfo) (Student’s t-test against β_CNxEI_ = 0, t(29) = −1.52, p > 0.1). The mean effect of the interaction on firing rates across the whole population was significant and negative, as 71 (57%) cells showed a negative influence and 53 (43%) a positive influence, with a mean regression weight (β_CNxEI_) = −0.056 ± 0.025 (spikes/sec)/(choice*bits_info_) (Student’s t-test, t(123) = −2.24, p < 0.05). As the trial progressed, PCC neurons generally showed lower firing rates for more informative expectations, matching the decrease in subject response times and in choice variability for larger information expectations. The full results of our GLM are reported in table S10.

**Table S10:** Full numbers of cells encoding all covariates for anticipation epoch, all choices inclusive (including diverge choices).

| Covariate | Monkey L (84 neurons) | Monkey R (40 neurons) |
| --- | --- | --- |
| Choice number (CN) | 15 | 2 |
| Expected reward (ER) | 4 | 0 |
| Expected information (EI) | 18 | 3 |
| Previous choice info (I-1) | 13 | 3 |
| Previous choice reward (R-1) | 17 | 4 |
| CN x ER | 4 | 0 |
| CN x EI | 23 | 7 |
| CN x I-1 | 8 | 3 |
| CN x R-1 | 11 | 2 |
| ER x EI | 11 | 2 |
| ER x I-1 | 15 | 1 |
| ER x R-1 | 0 | 0 |
| EI x I-1 | 22 | 5 |
| EI x R-1 | 15 | 0 |
| I-1 x R-1 | 9 | 2 |

### Additional Behavioral Entropy Encoding Results by Monkey

In the anticipation epoch, 51 neurons in monkey L (37 β’s > 0, 14 β’s ≤ 0; mean β_BE_ = 0.0077 ± 0.0045, t(50) = 1.70, p = 0.095; mean across all 84 neurons β_BE_ = 0.0061 ± 0.0028, t(83) = 2.15, p < 0.05) and 19 neurons in monkey R (14 β’s > 0, 5 β’s ≤ 0; mean β_BE_ = 0.0062 ± 0.0033, t(18) = 1.86, p = 0.078; mean across all 40 neurons β_BE_ = 0.0051 ± 0.0014, t(39) = 2.45, p < 0.05) significantly predicted behavioral entropy.

### Additional Encoding Results of Time of Receipt of Last Information

In the main text, we reported that 98 of 124 neurons distinguished before and after receiving the last information. This fraction was similar across the two monkeys: monkey L: 67 neurons (80%); monkey R: 31 neurons (78%).

### Additional Behavioral Entropy Encoding Results Before and After Receiving Last Information

Each trial’s binned spike counts were regressed against the time in the trial (TT), time of last informative outcome (TL), and their interaction (TTxTL). Of the cells significantly encoding the different states, 21 (21%) fired less and 77 (79%) fired more after the animal received all information compared to before, with a mean regression coefficient β_TTxTL_ = 0.017 ± 0.0030 spikes/information period (Student’s t-test against β_TTxTL_ = 0, t(97) = 5.52, p < 5 x 10’^−7^). Across the entire population, 30 (24%) fired more and 94 (76%) fired less before compared to after, with a mean regression coefficient β_TTxTL_ = 0.014 ± 0.0025 spikes/information period (Student’s t-test, t(123) = 5.52, p < 5 x 10^−7^).

Higher firing in individual neurons tended to predict behavioral entropy both before and after receiving the last bit of information. We next examined the size of the modulation of this choice-by-choice signaling. In our population, 74 (60%) cells predicted behavioral entropy before receiving the last informative outcome (all cells: mean β_BE_before_ = 0.0016 ± 0.00064 bits/spike, Student’s t-test: t(123) = 2.43, p < 0.05; significant cells: mean β_BE_before_ = 0.0018 ± 0.00096 bits/spike, Student’s t-test: t(73) = 1.83, p = 0.071) whereas only 24 (19%) predicted behavioral entropy after (all cells: mean β_BE_after_ = 0.0025 ± 0.0012 bits/spike, Student’s t-test: t(123) = 2.05, p < 0.05; significant cells: mean β_BE_after_ = 0.0044 ± 0.0022 bits/spike, Student’s t-test: t(23) = 2.27, p < 0.05; Fig. S3). Though the mean normalized population response failed to differentiate high from low average behavioral entropy trials after receipt of the last information as reported in the text, the choice-by-choice analysis of behavioral entropy revealed consistent and significant positive modulation both before and after. In sum, averaging the modulation of neuronal firing rates showed that PCC neurons showed higher firing for high behavioral entropy both before and after, albeit with many fewer significant cells after.

### Supplemental Figures

**Figure S1.**
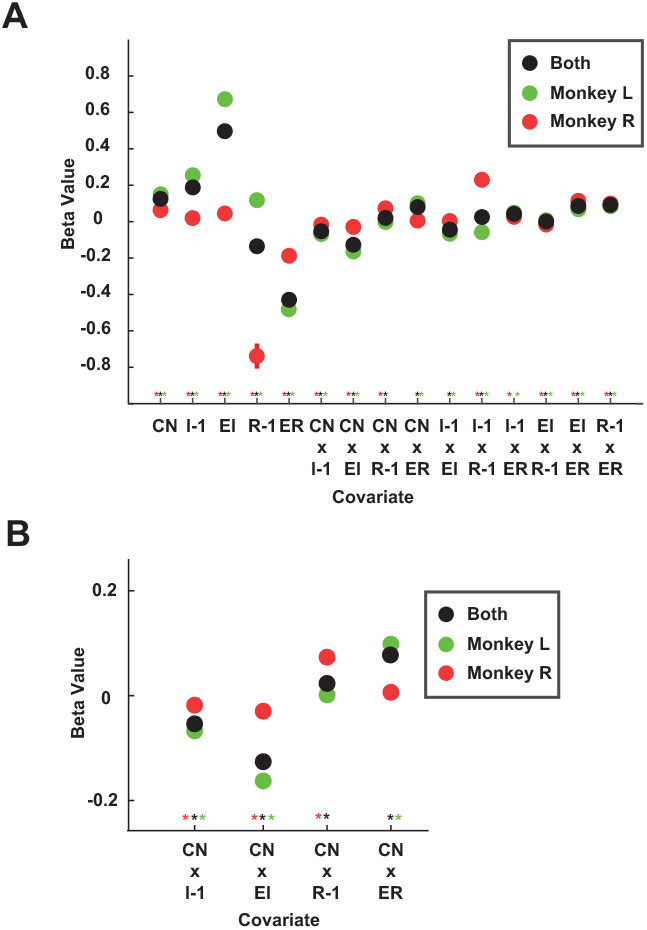
Coefficient values from response time and behavioral entropy regressions. **A**. coefficient values from regression of response time against listed covariates. **B**. coefficient values from regression of behavioral entropy against listed covariates. CN = choice number; EI = expected information; I – 1 = previous choice information outcome; ER = expected reward; R – 1 = previous choice reward outcome. Error bars either occluded by points or ± 1 s.e.m. * = p < 0.05, Bonferroni corrected.

**Figure S2.**
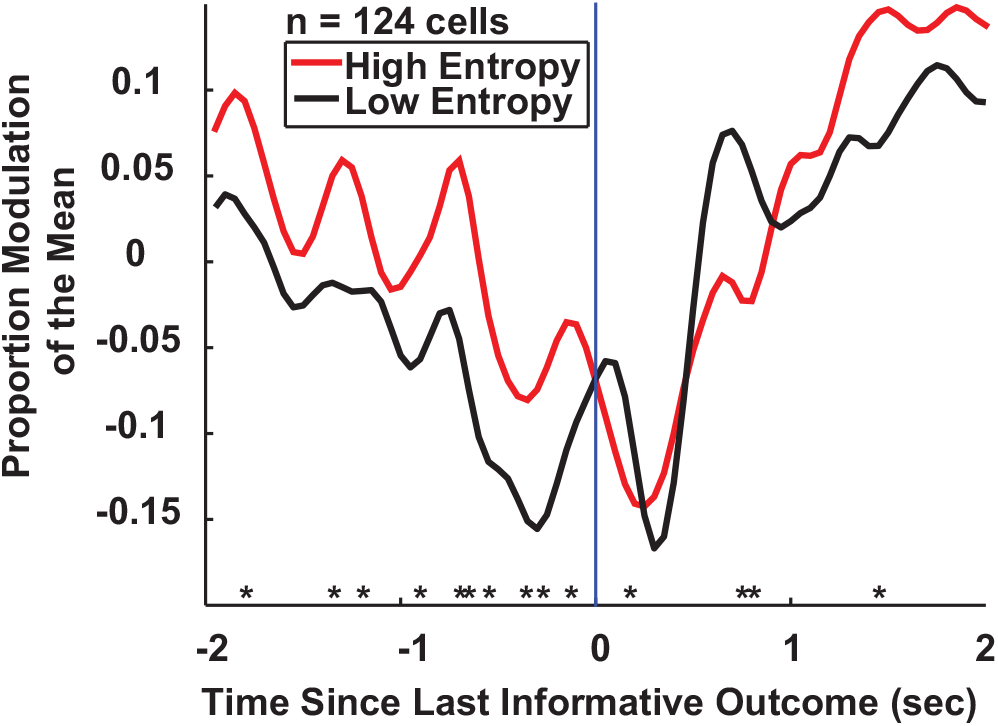
The multiplexing of behavioral entropy and time of receipt of last information in the PCC population. The population multiplexed information boundary signaling with behavioral entropy encoding, with higher firing rates for more variable choices before the last informative outcome but not after. n = 124 cells; * = p < 0.05; blue line = time of last informative outcome.

